# Subdivision of light signalling networks contributes to cellular partitioning of C_4_ photosynthesis in maize

**DOI:** 10.1101/465872

**Authors:** Ross-W. Hendron, Steven Kelly

## Abstract

Plants coordinate the expression of genes required to conduct photosynthesis in response to growth and environmental changes. In species that conduct two-cell C_4_ photosynthesis, the expression of photosynthesis genes is partitioned such that leaf mesophyll and vascular sheath cells accumulate different components of the photosynthetic pathway. The identity of the regulatory networks that facilitate this partitioning are unknown. Here we show that differences in light perception between mesophyll and bundle sheath cells facilitate differential regulation and accumulation of photosynthesis genes in the C_4_ crop *Zea mays* (maize). We show that transcripts encoding photoreceptors differentially accumulate in mesophyll and bundle sheath cells in a manner that is consistent with biophysical light filtration. We further show the blue light (but not red) is necessary and sufficient to activate photosystem II assembly in etiolated maize mesophyll cells, while both red and blue produce the same effect in C_3_ *Hordeum vulgare* (barley). Finally, we demonstrate that changes in abundance of >20% of genes that differentially accumulate between mesophyll and bundle sheath cells can be recapitulated by spectrum specific deetiolation of maize seedlings. These findings provide evidence that subdivision of light signalling networks is a key component of cellular partitioning of C_4_ photosynthesis in maize.

## Introduction

Light is of fundamental importance to photoautotrophs, who harness energy from photons to synthesise sugars. Given the central role that light plays in the growth of plants it is fitting that they have evolved sophisticated methods to sense it and use this environmental cue to optimise their photosynthetic capabilities (Jiao et al., 2007). In the model plant *Arabidopsis thaliana* five families of photoreceptors, of varying spectral sensitivities, allow discrimination between different wavelengths of light, these comprise the phytochromes, cryptochromes, phototropins, UVRs and Zeitlupe proteins (Rizzini et al., 2011; Lin, 2000; Christie et al., 2015; Smith, 2000; Schepens et al., 2004). In brief, red light stimulates phytochrome signalling, blue light stimulates cryptochromes, phototropins and Zeitlupes, while UV-B stimulates UVR8. These photoreceptors either activate signalling cascades (Petroutsos et al., 2016), bind DNA directly (Yang et al., 2018), or bind to and affect the DNA binding properties of many transcription factors that serve as light signalling intermediates (Li et al., 2011; Yu et al., 2010). By linking light cues to the regulation of photosynthesis genes, plants are able to coordinate development of chloroplasts and optimise photosynthetic rates in mature leaf tissue under changing light conditions (Waters & Langdale, 2009; Ohashi-Kaneko et al., 2005; Ort et al., 2011).

In most angiosperm species, photosynthesis occurs in the leaf mesophyll. In dicotyledonous species, this cell layer contains at least two distinct cell-types; the palisade cells that are located adjacent to the upper leaf epidermis and the spongy mesophyll cells located beneath them. All light that strikes a leaf surface is filtered as it penetrates through these cell layers, such that light availability in the spongy mesophyll is reduced in comparison to the palisade by as much as 90% (Cui, Vogelkann & Smith, 1991; reviewed in Terashima et al., 2009). This biophysical light filtration is responsible for a decreasing gradient in chlorophyll and photosynthetic capacity through the mesophyll such that upper palisade cells carry out vast majority of photosynthesis in the leaf (Evans & Vogelmann, 2003; reviewed in Tholen et al., 2012). In addition, different wavelengths of photosynthetically active light vary in their ability to penetrate leaf tissues, with longer wavelength red light penetrating significantly deeper than shorter wavelength blue light (Vogelmann & Evans, 2002; Slattery et al., 2016). Modelling light penetration through the leaf suggests that just 4% of incoming diffuse blue light reaches the second layer of cells. In contrast, 12% of red light reaches these deeper cells, and thus both the intensity and spectrum of light change as light passes through the leaf (Xiao et al., 2016). Although monocotyledonous leaves contain more uniformly shaped mesophyll cells, analogous light filtration is thought to occur (Takahashi et al., 1994; Kume et al., 2017).

While most plant species conduct all the reactions required to carry out photosynthesis in a single cell, some species have evolved a way to split the process between two specialized cell types. This spatial division of photosynthesis is known as C_4_ photosynthesis, and it has evolved independently at least 61 times in both eudicots and monocots (Sage et al., 2016). The result of this spatial partitioning is a depletion of O_2_ and an increase in CO_2_ around RuBisCo that together reduce energy loss through photorespiration (von Caemmerer & Furbank, 2016). In all two-cell C_4_ species, these two specialised cell types are arranged concentrically in an outer layer of photosynthetic carbon assimilation tissue (PCA) surrounding an inner layer of photosynthetic carbon reduction tissue (PCR), which in turn typically surrounds the vascular tissue (Nelson & Langdale, 1989; Muhaidat, Sage & Dengler, 2007). This concentric wreath-like arrangement is known as Kranz anatomy (Haberlandt 1884). Thus, in two-cell C_4_ species all light reaching the PCR cell layer must first pass through an outer PCA cell layer. This means that in two-cell C_4_ species with Kranz anatomy, light reaching the PCR layer is ~10 fold dimmer and depleted in blue relative to red wavelengths compared to light that reaches the PCA layer. This light filtering is thought to be one reason why photosynthetic rates are lower under blue light than red in C_4_ species such as *Zea mays* and *Miscanthus x giganteus* (Sun et al., 2012, Sun et al., 2014).

Given the integration of light signals with the transcriptional control of photosynthesis (reviewed in Wang, Hendron & Kelly, 2017), it was hypothesised that photoreceptors may differentially accumulate between PCA and PCR cells in a manner that is consistent with biophysical light filtration, and may thus play a role in facilitating partitioning of photosynthetic reactions in C_4_ species. To test this hypothesis the differential transcript accumulation of photoreceptors was investigated in the PCR (in this case bundle sheath) and PCA (in this case mesophyll) cells of mature leaves of C_4_ plants representing independent origins of C_4_ photosynthesis. This revealed conserved partitioning of light regulatory networks, with blue light photoreceptors more prominent in mesophyll cells and red light equivalents more prominent in bundle sheath cells. This led us to further hypothesise that gene regulatory networks underpinning chloroplast development in mesophyll cells could be selectively stimulated by blue light. This was tested by exposing etiolated maize seedlings to blue or red light, and revealed that blue light but not red resulted in rapid accumulation of chlorophyll fluorescence consistent with functional PSII assembly. Consistent with the partitioning of photoreceptor gene expression and light availability within the leaf, transcripts encoding mesophyll cell specific genes were activated by light, while transcripts encoding bundle sheath specific genes were less light responsive. Moreover, while transcripts encoding mesophyll cell specific genes increased in abundance in response to either red or blue light, transcripts encoding bundle sheath specific genes increased in abundance in response to red light. Together these findings provide evidence that subdivision of light signalling networks contributes to cellular partitioning of C_4_ photosynthesis in maize.

## Results

### The cellular distribution of photoreceptors is consistent with relative light availability among cell types

To facilitate comparative analysis of the differential cell type accumulation of transcripts corresponding to photoreceptors in C_4_ grasses, orthologs of the experimentally validated *Arabidopsis thaliana* photoreceptor genes were identified in *Setaria italica, Sorghum bicolor* and *Zea ma*ys (Supplemental File S1). RNA-seq datasets for *S. viridis* (John et al., 2014), *S. bicolor* (Emms et al., 2016) and *Z. mays* (Chang et al., 2012) were used to quantify transcripts abundances and identify those that differentially accumulate between mature leaf mesophyll cells and bundle sheath strands in the three species (Supplemental File S1). The longest wavelengths of light that plants can detect are red/far red. These wavelengths are sensed by the phytochrome family of photoreceptors, of which there are five members in *A. thaliana*, PHYA to PHYE, and just three in grasses (Mathews & Sharrock, 1996). Consistent with the increased ratio of red light to blue light in bundle sheath cells, transcripts encoding 7 of the 12 phytochromes present in these species set exhibit significant preferential accumulation in bundle sheath cells while none exhibit preferential mesophyll cell expression (Figure 1A).

**Figure 1.**
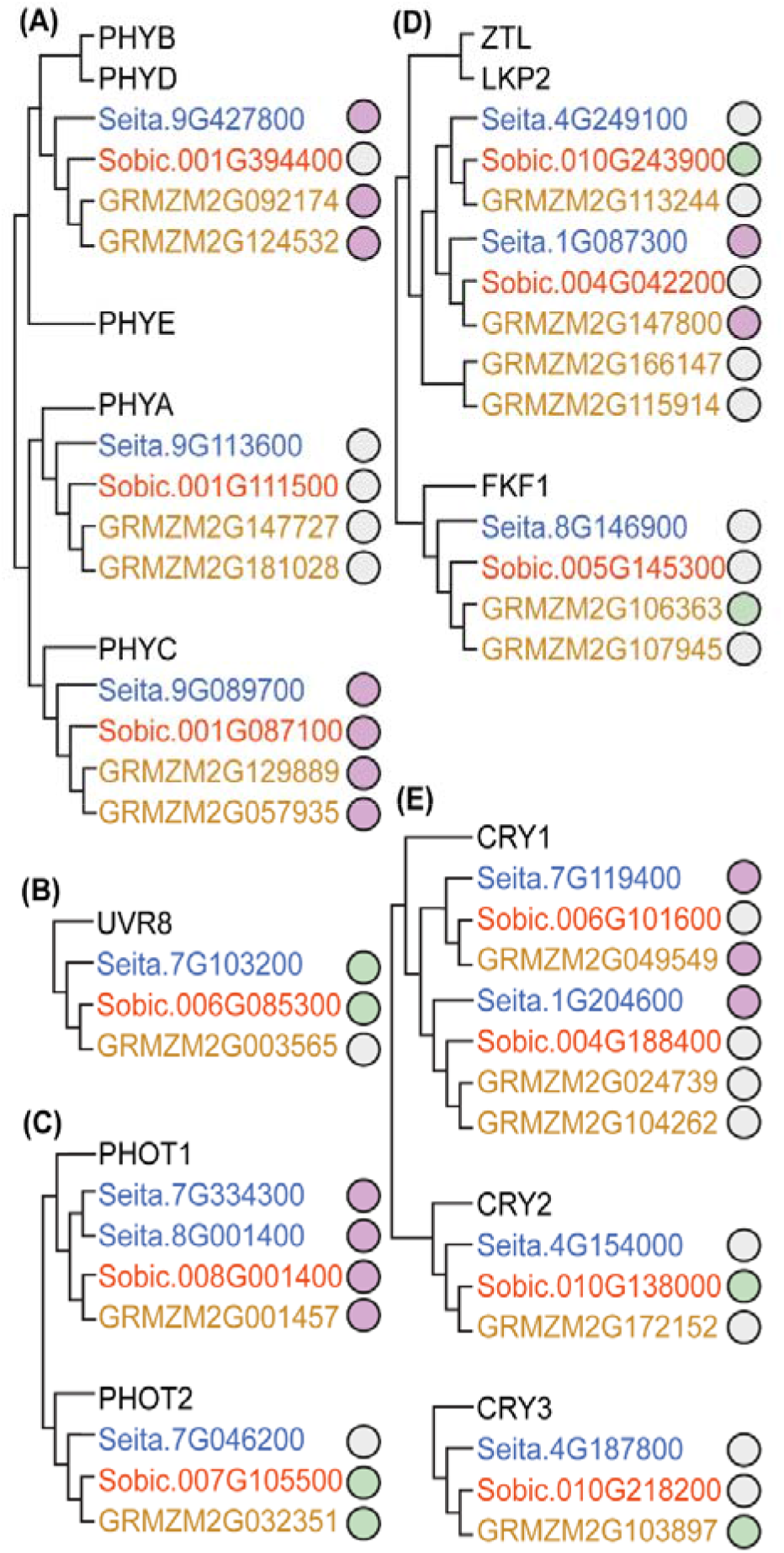
Differential accumulation of grass photoreceptors orthologous to A. thaliana (A) PHYs, (B) UVR8, (C) PHOTs, (D) ZTLs, and (E) CRYs in mesophyll and bundle sheath cells. *A. thaliana* common names label related grass clades, encompassing all *S. italica, S. bicolor* and *Z. may* ortholog*s*. Evolutionary relationships are indicated by a cladogram (left of each subplot). A purple dot indicates significant bundle sheath association, green for mesophyll association, and grey for not significant differential accumulation (at FDR < 0.01).

In contrast to red light, Ultra Violet B radiation is the shortest wavelength of light that plants can sense and in *A. thaliana* these wavelengths have been shown to mostly be filtered out in the top layer of photosynthetically active cells in the leaf (Bernula et al., 2017). In C_4_ species with Kranz anatomy this top layer is comprised of mesophyll cells and thus relatively little UVB should be available for perception in deeper cell layers. Consistent with the light availability in C_4_ leaves, the two orthologs of the UVB photoreceptor (UVR8) accumulate preferentially in mesophyll cells in sorghum and maize (Figure 1B).

The wavelengths of blue light are sensed by a larger complement of photoreceptors; the phototropins (Figure 1C), Zeitlupes (Figure 1D) and cryptochromes (Figure 1E). Within these gene families there are examples of both mesophyll and bundle sheath preferential accumulation of individual genes. The clearest example of spectral partitioning occurs for the phototropin (PHOT) genes. All four PHOT1 transcripts preferentially accumulate in the bundle sheath, whilst the maize and sorghum PHOT2 orthologs preferentially accumulate in the mesophyll. This partitioning is noteworthy because a key difference between these two clades of PHOT genes is their sensitivity to blue light intensity. PHOT2 mediates phototropism and chloroplast avoidance mechanisms under high intensity blue light whilst PHOT1 is associated with low intensity blue light (Kasahara et al., 2002). Thus, the cell type partitioning of PHOTs in C_4_ species matches the sensitivity of the photoreceptor to the intensity of the incident blue light in each cell type (higher intensity in mesophyll, lower intensity in bundle sheath). Another key difference is that PHOT2, but not PHOT1, mediates the chloroplast avoidance response (Gotoh et al., 2018). This is pertinent as PHOT2 is expressed in C_4_ mesophyll cells in which chloroplasts exhibit the avoidance response (Maai, Miyake & Taniguchi, 2011), while PHOT1 is expressed in the bundle sheath in which chloroplasts do not exhibit the avoidance repose (Maai, Miyake & Taniguchi, 2011). Thus, partitioning of PHOT genes not only matches the expression of the photoreceptor to the light intensity that is available, but may also explain the chloroplast behavioural differences between the two cell types.

Likewise, CRY1 differs from the constitutively nuclear localised CRY2 and CRY3 in that it is rendered inactive when exposed to blue light via export from the nucleus (Lin & Shalitin, 2003). It is therefore fitting that all three differentially accumulating CRY1 genes are preferentially expressed in the bundle sheath, where blue light is a weaker signal, whilst the differentially expressed CRY2 and CRY3 orthologs instead accumulate in the mesophyll (Figure 1E), where blue light can maximally stimulate their activity.

In total, for the 11 C_4_ monocot photoreceptor gene families (related to PHYB/PHYD, PHYA, PHYC, UVR8, PHOT1, PHOT2, ZTL/LKP2, FKF1, CRY1, CRY2 and CRY3) maize exhibits preferential bundle sheath or mesophyll cell transcript accumulation of at least one copy of 8 of these photoreceptors, *Setaria* 6 and *Sorghum* 6. Thus, the photoreceptors that initiate light signalling networks show biased cell type accumulation patterns that are generally consistent with the properties of the incident light that those cell types receive. Moreover, within this sample, the partitioning of photoreceptors between cell types appears to be most pronounced in maize.

### Blue light, but not red, stimulates assembly of photosystem II in etiolated maize seedlings

Given that transcripts encoding photoreceptors displayed differences in their cell-type accumulation patterns that was consistent the with light spectrum available to those cell types, it was hypothesised that maize may have exploited this phenomenon to differentially partition the expression of downstream photosynthesis genes between bundle sheath and mesophyll cells. To test this hypothesis, we exploited the fact that many C_4_ monocots and dicots exhibit preferential accumulation of photosystem II (PSII) in mesophyll cells through both transcriptional and post-translational mechanisms (Meierhoff & Westhoff, 1993, Hofer et al., 1992). This preferential accumulation must be mediated by photosynthesis gene regulatory networks that are active in the mesophyll and inactive in the bundle sheath. Furthermore, maize preferentially accumulates blue light sensitive photoreceptors in the mesophyll (Figure 1) and assembles PSII only in the mesophyll and not the bundle sheath due to light-dependent transcription in mesophyll cells (Schuster et al., 1985, Sheen at al., 1988). Hence, it was posited that activation of blue light signalling networks would specifically enhance PSII assembly in maize while activation of red light signalling networks would not. In contrast, C_3_ species should not exhibit this partitioning as both light signalling networks are active in the same cell type and thus either blue or red light should activate PSII expression and assembly.

To test this hypothesis, maize seedlings were de-etiolated by illumination with either 100 μmol m^−2^ s^−1^ blue or red light and functional PSII assembly was monitored by chlorophyll fluorescence after two hours. This revealed significant PSII assembly under blue light but none under red light (Figure 2A). In order to generate a time series for PSII assembly this experiment was expanded upon by taking measurements every 15 minutes for three hours for both maize and barley seedlings. In barley, either red or blue light alone were sufficient to induce rapid increases in фPSII (Figure 2B). This is compatible with previous findings in barley where PSII protein and water splitting were detectable within one hour after exposure to 30 μmol m^−2^s^−1^ of white light (Shevela et al., 2016). In contrast, in maize analogous induction of functional PSII was only stimulated by blue light (Figure 2C), with only a minor increase in фPSII after three hours of stimulation with red light (Figure 2C). Thus, the light signalling network governing фPSII development in barley was equally receptive to blue and red light but in maize the mesophyll regulatory network has reduced red light sensitivity and was strongly activated by blue light signalling.

**Figure 2.**
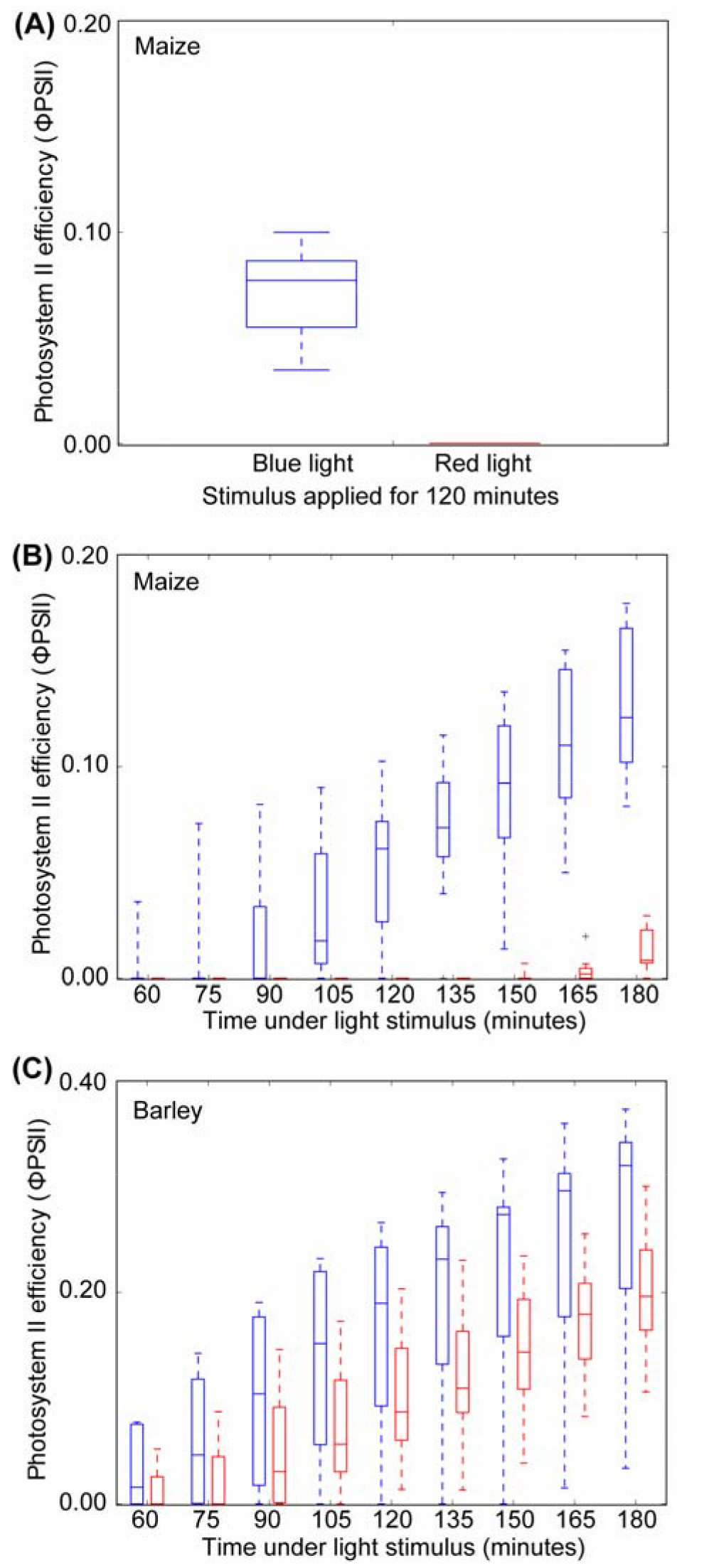
Grouped box plots indicating фPSII as a function of time under red (red boxes) or blue (blue boxes) light during de-etiolation with 95% confidence intervals displayed. (A) Maize leaves measured once after two hours (n=5), (B) de-etiolation time series of maize leaves (n = 9), (C) de-etiolation time series of barley leaves (n = 9).

### Activation of blue light signalling networks promotes the transcription of PSII assembly components

Whilst the rapid increase in PSII under blue light was predominantly due to post-translational assembly of PSII monomers already in the etioplast (Muller & Eichacker, 1999; Forger & Bogorad, 1973), it was also likely to be driven by spectrum specific transcriptional responses. To investigate this, total RNA was isolated from leaves at the end of the red or blue light induction period above and subject to transcriptome sequencing. Samples were also taken from etiolated leaves that received no light treatment. The complete set of replicated mRNA abundance estimates, differential testing results as well as gene accessions and TPMs are accessible in the Supplemental File S1. To simplify nomenclature, and for consistency with extant literature, *A. thaliana* homolog names are used throughout.

Given that blue light specifically induced the assembly of PSII in maize, it was hypothesised that the subunits of the PSII complex and its assembly factors would be upregulated under blue light. Of the seven subunits whose transcript abundance changed in response to light stimulation, three were preferentially upregulated under blue light (orthologs of PSBP, PSBE, and PSB28’, Figure 3A), while the other four preferentially responded to red light (PSBO, PSBQ, PSBY and PSB28’, Figure 3A). However, blue light preferential expression was observed for transcripts encoding the machinery that facilitates protein complex assembly in plastids, including TRANSLOCONS OF OUTER CHLOROPLAST (TOC) protein importers and molecular chaperone proteins such as HEAT SHOCK PROTEIN 90 (HSP90) proteins and CHAPERONIN 60 (CPN60A). Moreover, SEQUENCE RECOGNITION PARTICLE 54 (SRP54) and ALBINO 3 (ALB3), which insert the PSII light harvesting complex into thylakoid membranes (Shuenemann et al., 1998), and PSII turnover proteins (FtsH2 and FtsH5) (Kato et al., 2009) were all preferentially activated by blue light stimulation (Figure 3A). Thus, while the transcripts encoding components of the PSII complex were induced by both red and blue light, light responsive transcripts encoding PSII assembly factors were all preferentially induced by blue light. This suggests that the mechanism of PSII accumulation in the mesophyll of mature leaves is implemented at two levels – by blue light signalling networks acting on the post-transcriptional assembly of functional PSII complexes and by blue light signalling networks promoting the transcription of the requisite assembly factors.

**Figure 3.**
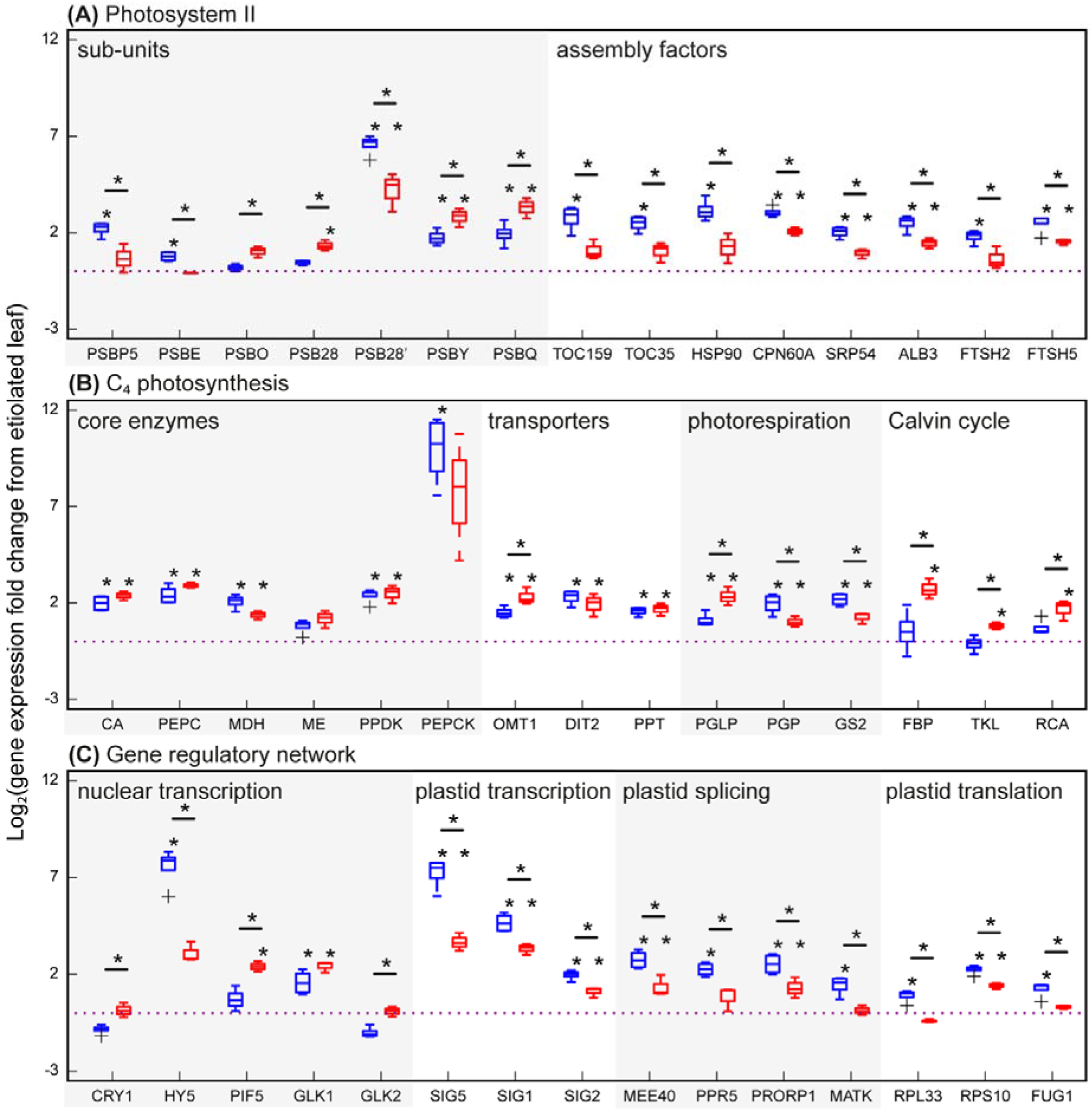
Differential expression of photosynthesis genes under blue and red light treatments. Asterisks above boxplots indicate differential expression between light treatment and leave kept in the dark, whilst asterisks between bars (over horizontal lines) indicate significant differential expression between light treatments (FDR < 0.05 and 95% confidence intervals shown). Dotted lines at y=0 indicate the average expression in etiolated tissue, from which fold changes were calculated. (A) Photosystem II subunits and assembly factors, (B) C_4_ carbon fixation machinery, including carbon concentration enzymes, transporters, photorespiration and Calvin cycle genes, (C) Regulators of gene expression relating to photosynthetic capacity.

### Transcripts encoding C_4_ cycle components are generally not differentially responsive to light spectrum

Previously it has been shown that transcripts encoding several proteins of the C_4_ cycle increase in abundance in response to light in a eudicot C_4_ species (Burgess et al., 2016). Given that blue light signalling networks promoted the development and expression of mesophyll specific PSII, it was hypothesized that light responsive components of the core C_4_ cycle may also exhibit light spectrum sensitivity. Consistent with previous analysis (Burgess et al., 2016), transcripts encoding C_4_ cycle enzymes and transporters in maize to increase in abundance in response to light (Figure 3B). However, in contrast to previous observations in the C4 eudicot *Gynandropsis (cleome) gynandra* (Burgess et al., 2016), light mediated induction of MALIC enzyme was not observed. This may be due to differences between the species or differences between the experimental design and treatment. Although, these transcripts increased in abundance in response to light, with the exception OMT1, they did not exhibit preferential activation in response to blue or red light stimulation (Figure 3B). Thus the transcription of core C_4_ cycle genes, while light responsive, were insensitive to light spectrum differences.

In contrast to C_4_ cycle components, transcripts encoding three components of the photorespiratory pathway did show differential accumulation under different light treatments: PHOTORESPIRATORY 2-PHOSPHOGLYCOLATE (PGLP), was preferentially induced by red light whilst paralogous PHOSPHOGLYCOLATE PHOSPHATASE (PGP), was preferentially induced by blue light, as was GLUTAMINE SYNTHETASE 2 (GS2). Additionally, transcripts encoding three enzymes for carbon reduction were differentially expressed and consistently red light preferential: Transcripts encoding two enzymes of the Calvin Benson Cycle increased in abundance specifically following red light induction-FBPASE (FBP) and TRANSKETOLASE (TKL) (Figure 3B). Furthermore, while transcripts encoding rubisco were not differentially affected by light, transcripts encoding RUBISCO ACTIVASE (RCA) were significantly upregulated in response to red light (Figure 3B). Thus although the components of the C_4_ cycle did not show spectrum specific responses, other pathway components that exhibit cell type preferential expression also exhibited spectrum specific induction.

### Blue and red light have contrasting effects on regulators of photosynthesis gene expression control

Given the observed differences in light responses for numerous photosynthetic components, it was hypothesised that the putative regulators of these components may also differentially accumulate in response to different light stimuli. Transcripts encoding several key regulators of nuclear encoded photomorphogenesis genes (as classified in Wang, Hendron & Kelly, 2017) showed different accumulation responses to light spectra. HYPOCOTYL ELONGATED 5 (HY5), a positive regulator of greening, showed significant induction in response to blue light, while in contrast several PHYTOCHROME INTERACTING FACTORS (PIFs) which are typically negative regulators of greening, showed red light activated expression (a PIF5 homolog is displayed in Figure 3C). Thus, the antagonistic transcriptional pathways that modulate greening promotion and inhibition appear to be sensitive to different wavelengths of light in maize. The GOLDEN-2 LIKE genes also respond differently to light treatments. GOLDEN2-LIKE 1 (GLK1) was activated equally in response to blue and red light stimulation, while GOLDEN2 (GLK2) expression was repressed by blue light and insensitive to red light (Figure 3C). This blue light mediated repression of GLK2 may provide a mechanistic explanation for the bundle sheath preferential expression of this gene in C_4_ grasses (Waters & Langdale, 2009, Wang et al., 2013). Of the photoreceptors, only transcripts encoding CRY1 accumulated differentially following red or blue light induction, with reduced abundance under blue light. Hence, blue light mediated gene networks induced negative feedback on this blue light photoreceptor.

It is critical that the gene expression machinery of the nucleus is coordinated with that of the plastid during greening. It is therefore noteworthy that all of the nuclear and plastid encoded components of the plastid transcription, splicing and translation machinery that increased in abundance in response to light were activated preferentially by blue light (Figure 3C). These included SIGMA factors SIG5, SIG1 and SIG2. Notably, 97 different pentatricopeptide repeat (PPR) containing proteins, increased in abundance following blue light illumination, while only three increased following red light (Supplemental File S1). The list of blue light induced PPR proteins includes several regulators of photomorphogenesis, including MATERNAL EFFECT EMBRYO ARREST 40 (MEE40) – which prevents delayed thylakoid development in rice (Su et al., 2012), PENTATRICOPEPTIDE REPEAT 5 (PRR5) – which prevents chloroplast ribosome deficiency in maize (Beick at al., 2008), and PROTEINACEOUS RNASE P 1 (PRORP1) – which prevents a pale green phenotype in *A. thaliana* (Zhou et al., 2015). This extensive induction of RNA editing proteins, in addition to the blue light induction of the chloroplast-encoded RNA editing protein, MATURASE K (MATK), indicates that RNA editing machinery was primarily activated by blue light. This blue light preference was also seen for numerous chloroplast ribosome components (e.g. RPL33 and RPS10) as well as translation initiation factor FU-GAERI 1 (FUG1) (Miura et al., 2007). Thus, in addition to the promoting the increase in abundance of transcriptional activators of photosynthesis associated genes, blue light stimulated the accumulation of multiple transcripts encoding proteins implicated in the enhancement of plastid protein production, with functions ranging from RNA editing to the translational control of photosynthesis machinery.

### Subdivision of blue and red light signalling networks between mesophyll and bundle sheath cells is a key component of cell type specification in maize

Given the findings above, it was hypothesised that there would be a transcriptome wide associated between the responsivity of transcript to light spectrum and the cell type preferential expression of that transcript in mesophyll and bundle sheath cells. Specifically, transcripts that accumulated in mesophyll cells would also be induced by blue light and transcripts that accumulated in bundle sheath would be preferentially induced by red light. To test this hypothesis, the set of genes whose transcripts were responsive to spectrum specific light stimuli were compared to the set of genes whose transcripts exhibited cell type preferential expression (Supplemental File S1). The majority of light induced genes (2,395 out of the total 3,458) were upregulated uniquely under blue light, with relatively few (160) being upregulated only under red light (Figure 4A). Overall, transcripts encoding ~25% of mesophyll genes increased in abundance in response to light while ~5% of bundle sheath genes exhibited the same effect (Figure 4B). While mesophyll genes were responsive to both red and blue stimulation (Figure 4C & D), bundle sheath genes were significantly less responsive to blue light and significantly more responsive to red than would be expected by chance (Figure 4C & D). Thus, the transcriptional networks that promote the accumulation of mesophyll genes and bundle sheath genes differ in terms of both light sensitivity and responses to specific light spectra. Overall, ~21% of the genes whose mRNAs differentially accumulate between mesophyll and bundle sheath cells showed light dependent transcriptional responses during deetiolation, and thus subdivision of light signalling networks between specialised C_4_ cell types is a key component of C_4_ photosynthesis partitioning in maize.

**Figure 4.**
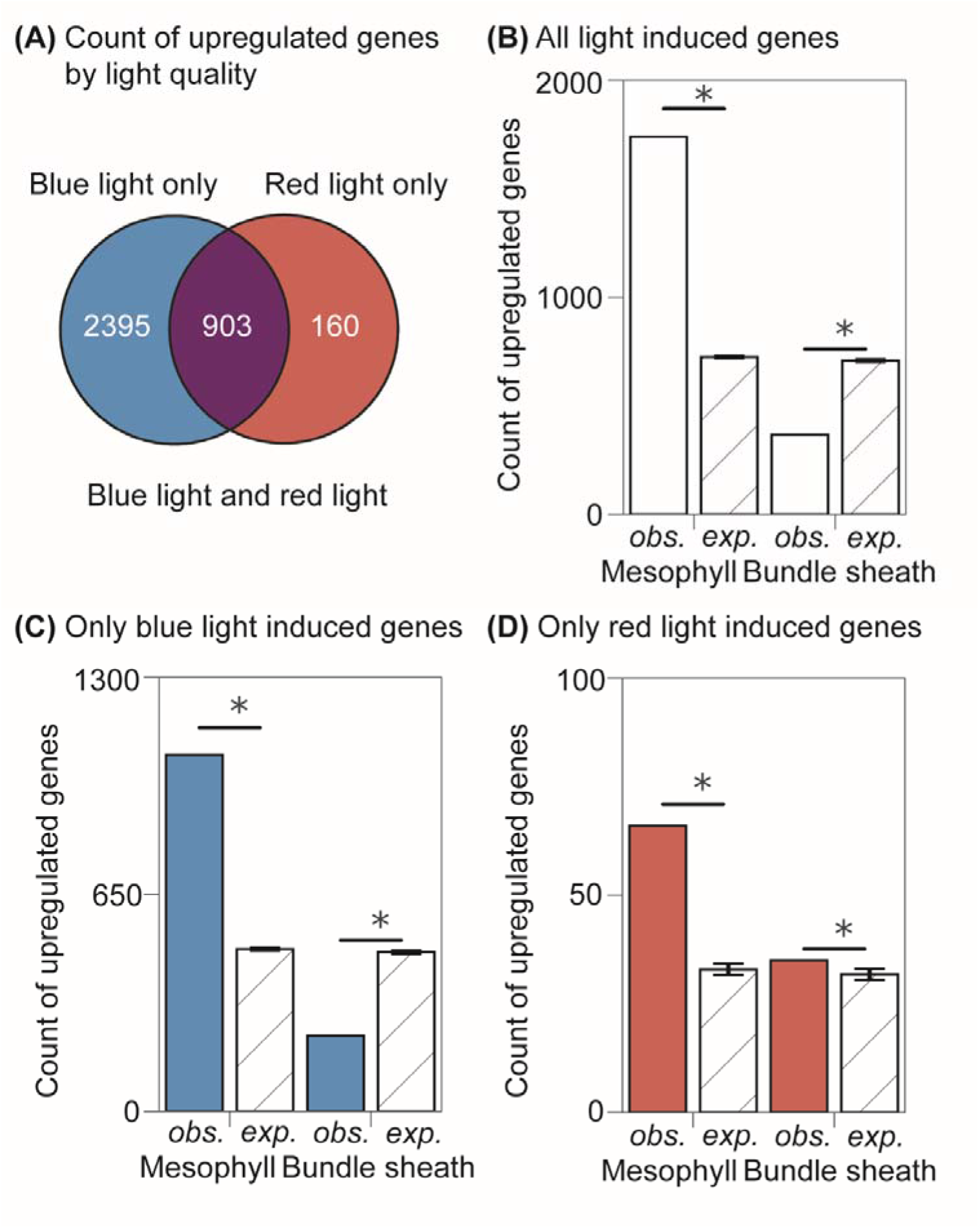
Association between light induction responses by mesophyll and bundle sheath genes. (A) The total counts of genes that were significantly upregulated by light during deetiolation, according to the type of light that induced them (red, blue or either). (B)-(D) Observed (obs.) and Expected (exp.) counts of mesophyll and bundle sheath genes that were induced by either type of light (B), only blue light (C) or only red light (D). Error bars indicate 99% confidence intervals for the mean expected counts generated by Monte Carlo resampling (n = 100) and asterisks indicate significantly different observations compared to null expectations at p < 0.01.

## Discussion

C_4_ photosynthesis is one of the most remarkable examples of anatomical, physiological and biochemical convergence in eukaryotic biology (Sage et al., 2016). The relative frequency with which it has evolved suggests that such large differences must be attributable to a small number of key regulatory changes (Wang et al., 2017). Here we show that blue and red light signalling networks are unevenly partitioned between mesophyll and bundle sheath in maize. Specifically, mesophyll cells preferentially accumulate components of light signalling networks while the bundle sheath accumulates fewer light signalling components but relatively more that are red light induced. Consistent with this partitioning we show that blue light, but not red light, is able to induce development of PSII fluorescence, a functional read-out of a protein complex that is specific to maize mesophyll cells. We further show that 21% of the transcripts that differentially accumulate between bundle sheath and mesophyll in mature maize showed altered transcript accumulation during de-etiolation, hence the partitioning of light networks accounts for a large component of cell type specific gene expression in maize.

Although, biased spectrum specific regulation of genes in the core C_4_ cycle was not observed, genes that encode other defining features of bundle sheath or mesophyll cells displayed biased spectral regulation. These included blue light responsive PSII assembly factors in the mesophyll and red light responsive photorespiratory pathway and Calvin Benson cycle components. Thus, biased sub-division of light signalling networks, while not directly responsible for cell-type specific patterns of transcript accumulation of core C_4_ cycle genes, can explain multiple distinguishing features of bundle sheath and mesophyll cells in maize. Together, these findings uncover a simple regulatory mechanism that facilitates photosynthetic partitioning between bundle sheath and mesophyll cells in maize.

Due to the extensive genetic redundancy prevalent in recently duplicated genomes such as that of maize (e.g. the majority of maize photoreceptors have paralogs, Figure 1), it will take considerable amount of time and experimentation to determine the full details of light signalling network partitioning between mesophyll and bundle sheath cells. In order to elucidate the molecular connections that link the photoreceptor profiles shown in Figure 1 to the downstream post-translational responses shown in Figure 2 and transcriptional responses shown in Figures 3 and 4, such work must be conducted at both transcriptional and post-transcriptional regulation levels in multiple null-mutant backgrounds. While knockouts of photoreceptor genes in maize remain elusive recent advances in genome engineering promise to accelerate this process.

Spectral filtering is a ubiquitous challenge for multicellular plants. Fittingly, a number of solutions have evolved to optimise light availability within the leaf. These include large scale solutions such as shade avoidance as well as developmental mechanisms driving dorsoventral asymmetry in photosynthetic capacity (reviewed in Terashima & Hikosaka, 1995). In all C_4_ species that exhibit Kranz anatomy the concentric arrangement of cell layers (Mudaidat, Sage & Dengler, 2007) inherently generates asymmetric light stimuli, such that the ratio of blue light to red light is higher in the photosynthetic carbon assimilation tissue than in the photosynthetic carbon reduction tissue. We propose that this purely biophysical phenomenon could represent a simple difference that became exploited during the evolutionary transition from C_3_ to C_4_ to help facilitate biased cell type expression of genes in C_4_ species (Figure 5). This model only requires that blue light signalling networks are more active in the outer cell layer than the inner cell layer, and therefore gene targets of blue light networks will preferentially accumulate in mesophyll cells. This may then be reinforced by the rewiring of gene regulatory networks such that, whilst mesophyll genes are enhanced by light, bundle sheath genes are more specifically enhanced by red light (Figure 4). Given the transcript abundance profiles of key photoreceptor genes in *Setaria viridis* (a representative species of an independent origin of C_4_ photosynthesis) it is likely that similar light mediated partitioning also occurs in this species and therefore it is tempting to speculate, that to some extent such a mechanism may be employed in other independent origins of C_4_ photosynthesis. Thus, the difference between mesophyll and bundle sheath cells in C_4_ species may be in part just a trick of the light.

**Figure 5.**
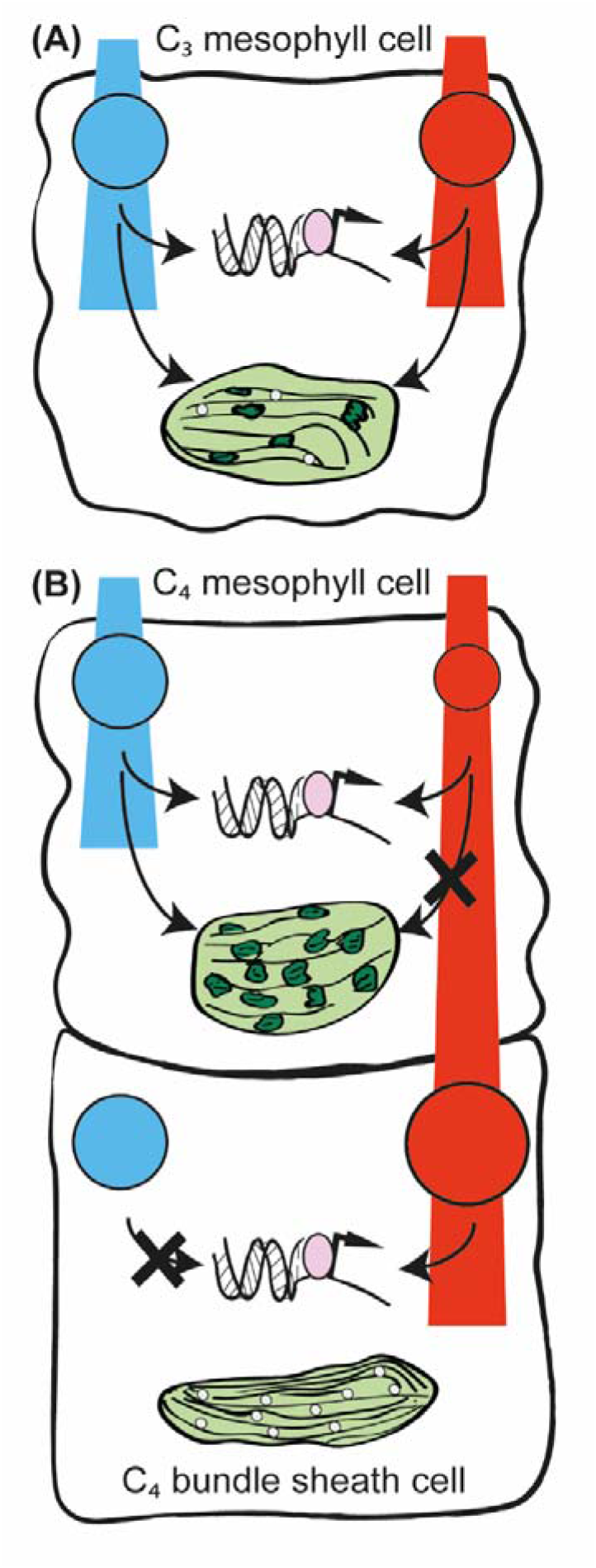
A model for C_4_ photosynthesis evolution from an ancestral C_3_ network. An ancestral network (A) of photoreceptors (displayed according to the colour of light they respond to and sized to indicate differential accumulation) regulate gene transcription and chloroplast development. Maturing photosynthetic chloroplasts contain thylakoids (stacked dark green bars) that synthesise starch (white dots). This partitions into a derived C_4_ network (B) whereby differential accumulation of photoreceptors, associated with the type of light reaching mesophyll and bundle sheath cells, fractures the downstream regulatory network. This alters regulatory interactions and results in two distinct chloroplasts, one with more thylakoid membranes, the other, synthesising starch. (B) Arrows summarise significant interactions presented in this study.

## Methods

### Identification of orthogroups and phylogenetic tree inference

The list of photosynthesis genetic regulators was compiled from Wang, Hendron & Kelly, 2017, for *Arabidopsis* and rice gene orthologs. Grass photoreceptor genes were identified by running Orthofinder V2.2.7 (Emms & Kelly, 2015) with setting ‘-S diamond’ on 16 plant species (Supplemental File S1), from which orthologs of known *A. thaliana* photoreceptors, and their phylogenies, were identified. Protein sequences were aligned using mafft-linsi (Katoh & Standley, 2013) and maximum likelihood gene trees were inferred using FastTree 2 (Price et al., 2010) for visual inspection of the photoreceptor phylogenies to check for any missed genes.

### De-etiolation of seedling leaves

B73 maize seeds were planted and grown in the dark for 9 days. Etiolated second leaves were clamped 5 cm from the leaf tip in a LICOR 6800 device equipped with a multi-phase flash fluorometer head. 100 μmolm^−2^s^−1^ of either 100% red or 100% blue light was used for de-etiolation. The LICOR was configured with flow rate 500 μmol s^−1^, 400 μmol mol^−1^ CO_2_, leaf temperature 28°C and 60% humidity. Fluorescence and gas exchange were measured either once after two hours or every 15 minutes for three hours. The consistency between both types of experiments at the two hour mark confirm that the measuring beam and fluorescent flashes (which use a red LED) were not providing a significant additional developmental stimulus when measurements were made more frequently. For barley, ‘Golden promise’ 7 day old seedling first leaves were sampled with the same settings. Negative фPSII values were assumed to be measurement artefacts of fully saturated reaction centres and so set to 0. Recently it has been shown that фPSII measurements can be erroneously inflated under blue light sources compared to red, resulting from over-estimation of the maximal fluorescence value (Evans et al., 2017). While this effect was shown to be not significant at the light intensity used in our experiment (Evans et al., 2017), it may have contributed to the difference in the absolute value of PSII observed between blue and red light induction in barley. It should be noted that this effect, even if present, cannot explain the lack of induction by red light in maize.

### RNA collection and sequencing

Maize leaf tissue that had either been exposed to blue light, red light or no light treatment were cut and frozen in liquid nitrogen immediately following the LICOR protocol. Tissue was ground and RNA extracted and purified using the Trizol method (Rio et al., 2010) in conjunction with a TurboDNAse treatment. Nine samples were sequenced using the BGIseq-500 RNA-Seq platform; four replicates of blue, three of red, two of no light treatment. The initial plan was to sequence three replicates of each, however one replicate of “no light” failed quality control prior to sequencing and thus the extra run was used to sequence an additional blue light sample.

### Analysis of gene expression data

Following sequencing, or fastq file download of raw read files, reads were trimmed using Trimmomatic version 0.38 (Bolger at al., 2014), with settings ‘LEADING: 10, TRAILING: 10, SLIDINGWINDOW:5:15, MINLEN:25’. Transcript counts and effective lengths were then quantified using Salmon version 0.10.0 (Patro et al., 2017) with settings ‘-l A - seqBias —gcBias’. Transcript counts from all gene models corresponding to the same gene were summed to generate abundance estimates at the gene locus level. For each species, raw counts for each locus were analysed using DEseq2 (Love et al., 2014), from which a set of maize mesophyll (6,873) and bundle sheath genes (6,723) were defined using an adjusted p value cut off q < 0.01 (Supplemental File S1). Mapman categories for maize genes were downloaded (Usadel et al., 2009) and hypergeometric testing was carried out to identify any functional groups that were enriched within the sets of differentially expressed genes during de-etiolation, with Bonferroni multiple test correction used throughout. The total population of genes whose expression could be detected across mesophyll and bundle sheath cells in maize (29,530) were subjected to Monte Carlo resampling (n = 100) to generate expected numbers of light induced mesophyll and bundle sheath genes. These expected counts were compared to the observed numbers of differentially expressed mesophyll and bundle sheath genes using 99% confidence intervals.

## Data availability

RNA-seq data for bundle sheath and mesophyll transcriptomes used in this study were previously published and available in NCBI SRA under the following accession numbers ERP004434 (*S. viridis*), ERP013053 (*S. bicolor*) and SRP009063 (*Z. mays*). The RNA-seq data generated in this study have been deposited to ArrayExpress under accession EMTAB-7200.

## Author Contributions

SK and RWH conceived the experiments. RWH conducted the analysis. SK and RWH wrote the paper.

## Acknowledgements

RWH is supported by a BBSRC studentship through BB/J014427/1. SK is a Royal Society University Research Fellow. This work was supported by the European Union’s Horizon 2020 research and innovation program under grant agreement number 637765 and by the BBSRC through BB/P003117/1.

